# Distribution, ecology, and natural history of the recently rediscovered and critically endangered Santa Marta Sabrewing

**DOI:** 10.1101/2024.03.18.581190

**Authors:** Esteban Botero-Delgadillo, Carlos Esteban Lara, Yurgen Vega, María Paula Santos, Diego Zárrate, Andrés M. Cuervo, John C. Mittermeier

## Abstract

The Santa Marta Sabrewing is a critically endangered (CR) hummingbird endemic to the Sierra Nevada de Santa Marta, Colombia. Prior to 2022, there were only three documented sightings of the sabrewing since it was described in 1879, including only one record between 1946 and 2022. As a result, this “lost” species has long been one of the most poorly-known birds in Colombia. We located a resident population of Santa Marta Sabrewing along the Guatapurí River near the Chemesquemena and Guatapurí villages in July 2022, and at its type locality, San José, in January 2023. Based on historical data and newly-collected field observations, we assess the species’ status and describe aspects of its natural history and ecology. Our review indicates the species has been frequently misidentified in the past, and that to date, documented evidence of its presence is limited to five localities, almost all of them restricted to the south-eastern slope of the Sierra Nevada de Santa Marta, along the mid Guatapurí river basin. Consequently, this bird appears to represent a case of microendemism. Following IUCN criteria, our data suggest that Santa Marta Sabrewing should remain listed as CR. Field observations indicate that the species is highly associated with watercourses, where males hold year-round territories and form leks. We obtained records of males in mid-elevation habitats (1,150–1,850 m) for 16 consecutive months between July 2022 and October 2023, suggesting that the species might not be an elevational migrant, as previously speculated. More information is needed to understand the species’ ecology so that effective conservation actions can be designed in collaboration with the indigenous communities with which the species coexists.

## Introduction

The Santa Marta Sabrewing (*Campylopterus phainopeplus*) is the rarest and most poorly-known of the 24 endemic bird species restricted to the Sierra Nevada de Santa Marta (SNSM), an isolated coastal mountain range in northern Colombia that is considered the most important continental centre of endemism in the world and Earth’s most irreplaceable nature reserve (Le Saout *et al*. 2013, Duran-Izquierdo and Olivero-Verbel 2021, BirdLife International 2023a). With only one documented record since 1946, this hummingbird is listed as critically endangered (CR) by the IUCN given its rarity and presumed very small population (< 50 mature individuals; BirdLife International 2023b). The lack of confirmed records also resulted in Santa Marta Sabrewing being listed as a “lost” species (Rutt and Mittermeier 2023) and included in the top 10 most wanted lost birds in 2021 by the Search for Lost Birds (https://www.searchforlostbirds.org).

The first specimens of Santa Marta Sabrewing were collected by English geographer and naturalist F. A. A. Simons during his expeditions to the south-eastern slope of the SNSM in 1878 (Simons 1879, 1881). The species was described by O. Salvin and F. D. Godman in 1879 based on Simons’ specimens (Salvin and Godman 1879). Simons spent seven months mainly in two localities named Atánquez and San José, now part of the Cesar Department, as well as in Alto de Macotama, now in La Guajira Department (Figure 1). Between 1945 and 1946, American ornithologist M. A. Carriker collected eleven additional specimens of Santa Marta Sabrewing for the National Museum of Natural History, Smithsonian Institution, Washington, D.C. One of those specimens was collected in San José, the type locality, while the remaining ten were collected in Chendúcua, located at a higher elevation along the Guatapurí basin (Figure 1). Since Carriker’s collection, there were no documented records of the sabrewing until 2010, when researchers took a photograph of a mist-netted male in El Dorado private reserve, Magdalena department, within the Important Bird Area Cuchilla de San Lorenzo, located ∼60–85 km west of the previously known localities (Figure 1).

**Figure 1.**
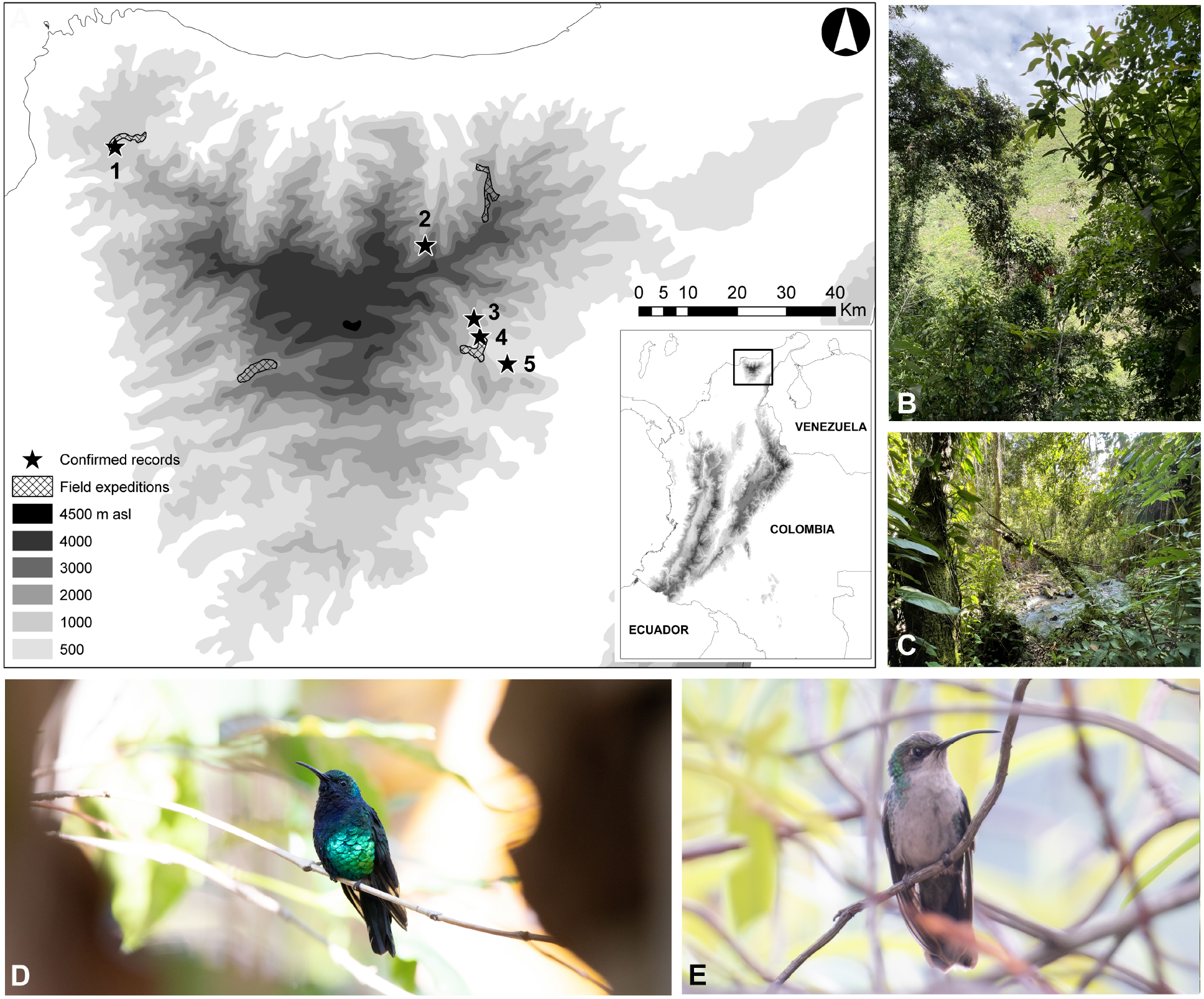
(A) Locations with confirmed records for Santa Marta Sabrewing in the Sierra Nevada de Santa Marta, northern Colombia, together with locations of field surveys conducted in 2022–2023. Numbered localities are: (1) San Lorenzo; (2) Alto de Macotama; (3) Chendúcua; (4) San José; (5) Atánquez (∼8 km from Chemesquemena). (B) The Guatapurí river basin has extensive areas of agricultural areas mixed with riparian forests, which are more common on steep slopes such as along La Macana stream near Chemesquemena. (C) Santa Marta Sabrewing was found in riparian forest strips nearby watercourses with dense understoreys. (D, E) Male and female sabrewings observed at the study site. Photographs by Elquin Toro, John C. Mittermeier, and Esteban Botero-Delgadillo © (reproduced with permission).

With such limited records, information on Santa Marta Sabrewing’s distribution, ecology and natural history is sparse. The species has been suspected to be an elevational migrant, with seasonal movements occurring between the dry and wet seasons (BirdLife International 2023b), although this has never been confirmed. While inhabiting humid forest borders and mixed landscapes of forest and shade-grown coffee plantations at 1,200–1,800 m asl during the dry season (February–May), it has been supposed to migrate to the paramo and up to the snowline at 4,800 m asl during the wet season (July–October; reviewed in Cárdenas-Ortiz and Cortés-Herrera 2016, BirdLife International 2023b). Literature on its natural history suggest that it consumes nectar from flowers of *Musa* and *Inga* trees inside coffee plantations and *Befaria* and *Palicourea* plants in forest edges, and that courtship displays take place from April to June (see Cárdenas-Ortiz and Cortés-Herrera 2016). However, there are no direct observations or any published evidence confirming foraging visits to coffee plantations or courtship behaviour. Most knowledge about the sabrewing’s ecology is based on anecdotal records, and the fact that it is regularly misidentified (see McMullan 2016) casts doubt on the validity of available information. For instance, males of the remarkably similar Lazuline Sabrewing (*Campylopterus falcatus*) and Black-throated Mango (*Anthracothorax nigricollis*) seem to be frequently misidentified as male Santa Marta Sabrewings, whereas females are confused with female White-vented Plumeteers (*Chalybura buffonii*). Some putative vocal recordings archived in acoustic online repositories have later been confirmed to be incorrectly assigned to the species (Salaman et al. 2022). Thus, it remains unclear whether the scarce information on Santa Marta Sabrewing is accurate and reliable.

Remarkably, a population of Santa Marta Sabrewing was located in July 2022 by YV near the upper part of the Chemesquemena village (Cesar Department; ∼8 km from Atánquez), not far from an area where Cristobal Navarro, a naturalist working on conservation projects with the Kogi indigenous people, may have seen the species in 2015 and 2019. In January 2023, CEL and AMC found isolated male individuals at a nearby locality in Chemesquemena and at San José (Figure 1). These findings were widely covered by national and international media and elicited a response from a consortium of research and conservation organizations to collect more information on the species. Here, we present the results of this effort including information collected over a 16-month field survey in four different localities in the SNSM (Figure 1). Specifically, we provide: (i) updated information on the species’ distribution from historical data and field expeditions conducted during 2022 and 2023; (ii) preliminary projections of the species’ extent of occurrence (EOO) and area of occupancy (AOO) based on confirmed presence records; and (iii) new information on the species’ ecology and natural history based on a monthly monitoring of the population at Chemesquemena, including local rate of occupancy, seasonal movements, habitat associations, social and breeding behaviour, and a first characterization of its vocal repertoire. We disccuss our findings in the context of the information currently available for the species and evaluate the conservation implications.

## Methods

### Geographic distribution assessment and field expeditions

We conducted a thorough revision of scientific literature, public databases, digital collections, and museum bird collections in search of confirmed records and specimens of Santa Marta Sabrewing. Public databases consulted included the Global Biodiversity Information Facility (GBIF; www.gbif.org), eBird (www.ebird.org), xeno-canto (www.xeno-canto.org), and national-level data repositories, including SiB (www.sibcolombia.net). Bird collections consulted were those with good representation of SNSM specimens, including Instituto Alexander von Humboldt (IAvH) and Instituto de Ciencias Naturales, Universidad Nacional de Colombia, in Colombia; the Carnegie Museum of Natural History, National Museum of Natural History (Smithsonian Instituton), and American Museum of Natural History in the United States; and the Natural History Museum in the United Kingdom. We only considered evidence-based records including photographs, videos, audio recordings, and specimens.

In addition to fieldwork around Chemesquemena and nearby areas, we also conducted 4– 7-day field expeditions between December 2022 and August 2023 to three other localities where Santa Marta Sabrewing could occur (Figure 1): the San Lorenzo IBA (Magdalena Department); the upper Rioancho river basin, in Dibulla (La Guajira Department), close to Alto de Macotama where there have been recent unconfirmed sightings (C. Navarro, pers. comm., [May 2022]); and Aracataca (Magdalena Department), a locality where the species could potentially occur according to recent distribution hypotheses.

We built a minimum convex polygon (MCP) based on the confirmed records of Santa Marta Sabrewing to project the species’ extent of occurrence (EOO). Subsequently, we projected the species’ area of occupancy (AOO) by overlapping the EOO polygon with a geographical layer of suitable habitat (see IUCN 2022). A suitable habitat layer was generated by selecting areas of mature and secondary forests, riparian forests, and areas of perennial crops mixed with patches of native vegetation (see BirdLife International 2023b), delimited by the extent of the three biomes where the species’s presence was confirmed (see *Results*). We compared our EOO and AOO projections with the threshold values given under the IUCN’s criteria B1 and B2 (IUCN 2022).

Suitable habitat delimitation was conducted in ArcGIS Desktop 10 (ESRI 2011) using a vegetation layer from the CORINE Land Cover Methodology for Colombia, 2018, at 1:100,000 scale (IDEAM 2021), and a layer of biomes compiled in the public-access portal of the Land-use Planning Geographic Information System (SIG-OT) of the “Agustín Codazzi” Geographic Institute of Colombia (https://geoportal.igac.gov.co).

### Population monitoring and occupancy

From September to December 2022, we conducted bird counts to estimate local occupancy and obtain a raw estimate of abundance along the course of La Macana stream (a tributary stream of the Guatapurí River) near Chemesquemena, where the sabrewing had been found by YV (hereafter referred to as the rediscovery site). Bird surveys were carried out using 20 georeferenced point-count stations (30 m radius), in which one observer (YV) searched for Santa Marta Sabrewing for 15 min and documented every visual or acoustic detection. One count per station was conducted every month between 07h00 to 11h00, with visits to each site alternating between early and midmorning across replicates. Count stations were located from 1,162 to 1,819 m asl (mean: 1,615 m) in a landscape dominated by four land cover types defined using the CORINE land cover layer: secondary vegetation (thickets and early succession vegetation of at least 2 m height); early growth (shrubberies and abandoned pastures covered with bushes and ferns); mixed vegetation (mosaics of pastures, native vegetation, and productive lands such as agroforestry); pastures and open areas. To estimate occupancy rate, we generated a detection history file consisting of all detection/non-detection data based on 80 surveys (four replicates at each of 20 point-count stations). Detection histories were analysed using the *occu* function in the *unmarked* package (Fiske and Chandler 2011) in R 4.0.2 (R Core Team 2020). Given the limited number of detections (*n* = 23), we fitted an intercept-only model to reduce the risk of over-parameterization. We then fitted two additional complex models to explore whether occupancy varied in response to elevation and/or land cover in the study area. One “saturated” model included as predictors the linear and quadratic effects of elevation and the proportion of native vegetation around each point-count (100 m buffer around the point location), whereas the second one was an “elevation-only” model. The proportion of native vegetation was estimated using a buffer analysis that allowed us to estimate the % of native vs. transformed vegetation around each point based on the CORINE land cover layer. The detection probability was held constant for all three models.

### Natural history and behavioural observations

We conducted *ad libitum* observations at the rediscovery site from July 2022 to October 2023 to describe different aspects of the species’ natural history. Over this period, YV made monthly 2–3-day visits to monitor individuals and territories, recording whether sabrewings were present along the course of La Macana stream (∼1,150–1,850 m). We qualitatively described feeding habits and aspects of social and breeding behaviour, including territorial and lekking displays, vocal activity, and lek conformation.

Habitat associations were evaluated by calculating the proportion of detections of Santa Marta Sabrewing obtained in each of the four land cover types registered during point counts (see above). We also investigated patterns of habitat selection, comparing the observed and expected frequencies of use for each land cover type. We estimated the proportional extension of all vegetation types using remote images and the CORINE Land Cover methodology for Colombia: secondary vegetation corresponded to 66% of all area surveyed; early growth to 5%; mixed vegetation to 25%; pastures to 4%. Habitat use proportions and the confidence intervals around them were then compared to expected proportions through a Fisher’s exact goodness-of-fit test. Expected values by habitat were obtained by dividing the total number of detections of Santa Marta Sabrewing by the proportion of the area covered by each habitat (i.e., the combined area of all point counts and a 100 m buffer around them). Neu’s selection ratio (the ratio of observed to expected frequencies; Neu et al. 1974) was also calculated for each land cover type.

### Vocal analyses

We conducted acoustic analyses to describe the sabrewing’s territorial call based on audio recordings of three different males obtained near Chemesquemena. We selected and analyzed a single high-quality cut per male of 20 seconds each. The first male (Male A) was recorded on 5 March 2023 at 09h00 at a small farm lot with coffee plantations intermixed with banana trees and patches of sugarcane (1,350 m asl, ∼150 m distance to La Macana stream). This bird was perched ∼4.5 m above the ground, and as no other individuals were present in the area, was likely the sole owner of its territory. The second male (Male B) was recorded on 21 May 2023 at 08h30 in an small secondary riparian forest patch, and was perched ∼2.5 m above the ground (1,480 m, ∼ 40 m distance to La Macana stream). This individual was at a lek site where 3–5 males were frequently observed interacting throughtout our study. During the recording of Male B, another male was simultaneously vocalizing nearby. The third male (Male C) was recorded on 27 May 2023 at 10h00 in a dense and relatively large patch of secondary premontane forest on a steep slope (1,780 m, ∼200 m distance to La Macana stream). This bird was perched at ∼3 m above the ground and while it was singing alone, a second male was observed ∼50 meters away. CEL obtained these recordings using a Zoom H6 recorder coupled with a Rode NTG4 phantom-powered shotgun microphone. The sampling rate was 48 kHz with a depth rate of 24 bits for Male A, and 44.1 kHz with a depth rate of 16 bits for males B and C. We visualized and analyzed the recordings using the Seewave package (Sueur et al. 2008) implemented in R.

## Results

### Geographic distribution

We found confirmed specimen records of Santa Marta Sabrewing from only four localities, all of them in or near the Guatapurí River basin: Atánquez, San José, and Chendúcua in Cesar Department; and Alto de Macotama in La Guajira Department (see Figure 1 and Supplementary Material Table S1). While Simons labelled his specimens as Atánquez, which is located in the Badillo River basin, some or all might have been collected nearby Chemesquemena in the Guatapurí basin (1,200 m; see Salvin and Godman 1879). In addition to specimen records, we found one photographic record of a male Santa Marta Sabrewing in the San Lorenzo IBA (Magdalena Department) from 2010.

Misidentified records have been excluded from public databases such as eBird or xeno-canto in previous years, but erroneous entries still need to be corrected or removed from other datasets. For instance, four female specimens collected in 2015 and deposited in the IAvH Ornithological Collection were rexamined and identified as female White-vented Plumeteers (G. Bravo, pers. comm., [May 2023]), but these records still are labelled as “Santa Marta Sabrewing” in the GBIF and SiB portals. Likewise, a sound recording that was originally attributed to a male Santa Marta Sabrewing (XC235605) from the San Lorenzo IBA in 2007, was an individual not seen well and hence not visually confirmed. We were able to confirm that those vocalizations corresponded to Black-throated Mango (https://xeno-canto.org/forum/topic/54177). There are also 16 listed occurrences dating from 2004 to 2017 in GBIF that are likely erroneous, being either linked to human observations without supporting documentation or to collected tissues that need to be reexamined such as those described above.

We did not find Santa Marta Sabrewing during our expeditions to San Lorenzo, Rioancho, or Aracataca in 2022–2023. This left the four historically reported localities and the San Lorenzo photograph as the only georeferenced records with documenting evidence. Based on these data, we projected the species’ EOO to be ∼586 km^2^ (Figure 2).

**Figure 2.**
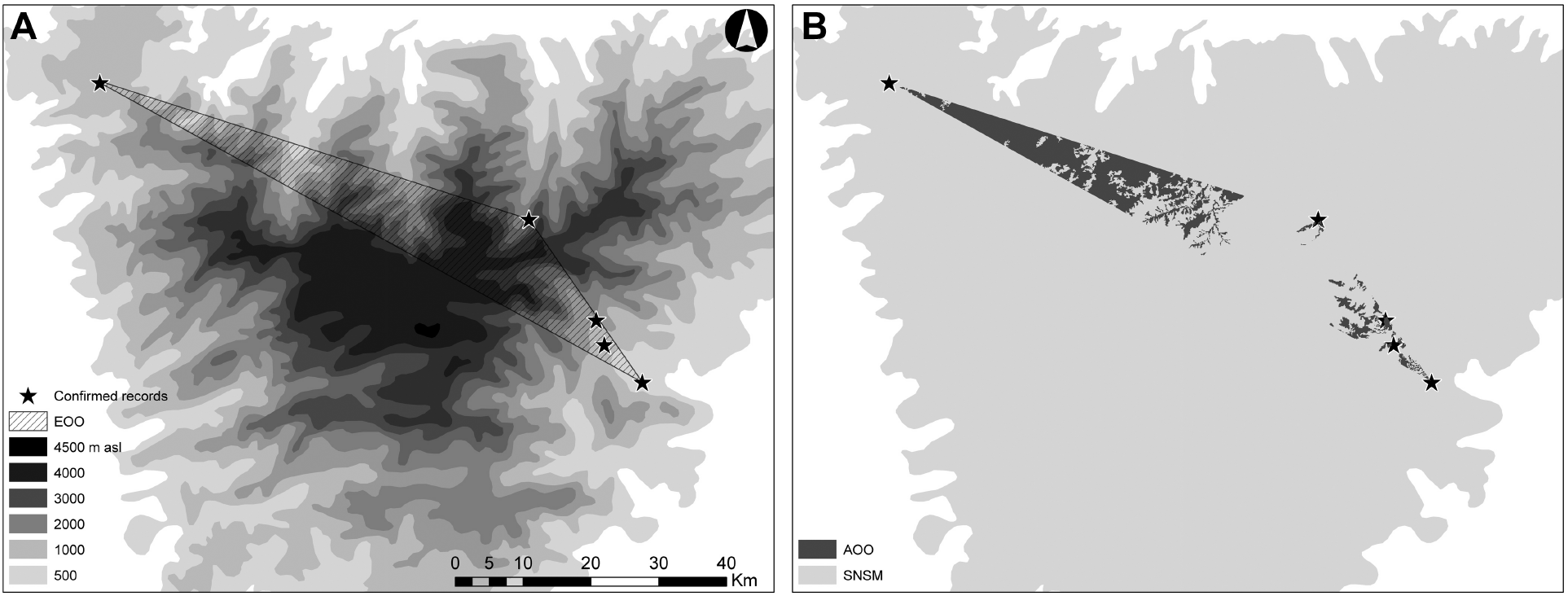
Spatial representation of Santa Marta Sabrewing’s (A) extent of occurrence (EOO), and (B) area of occupancy (AOO) in the Sierra Nevada de Santa Marta, northern Colombia.

Confirmed records of Santa Marta Sabrewing did not exceed elevations above 3,000 m, neither the exact coordinates nor a 2 km buffer around each point locality. Consequently, its documented presence was associated with three biomes: the Santa Marta and Macuira low orobiome; the Santa Marta orobiome; and the Santa Marta high orobiome. After intersecting areas of suitable habitat with these biomes within the species’ EOO, the AOO was projected to be ∼205 km^2^ (Figure 2). If the San Lorenzo record was excluded from this projection, as it could represent a case of vagrancy, the species’ AOO became restricted to the Guatapurí basin and covered ∼23 km^2^ (Figure 3A).

**Figure 3.**
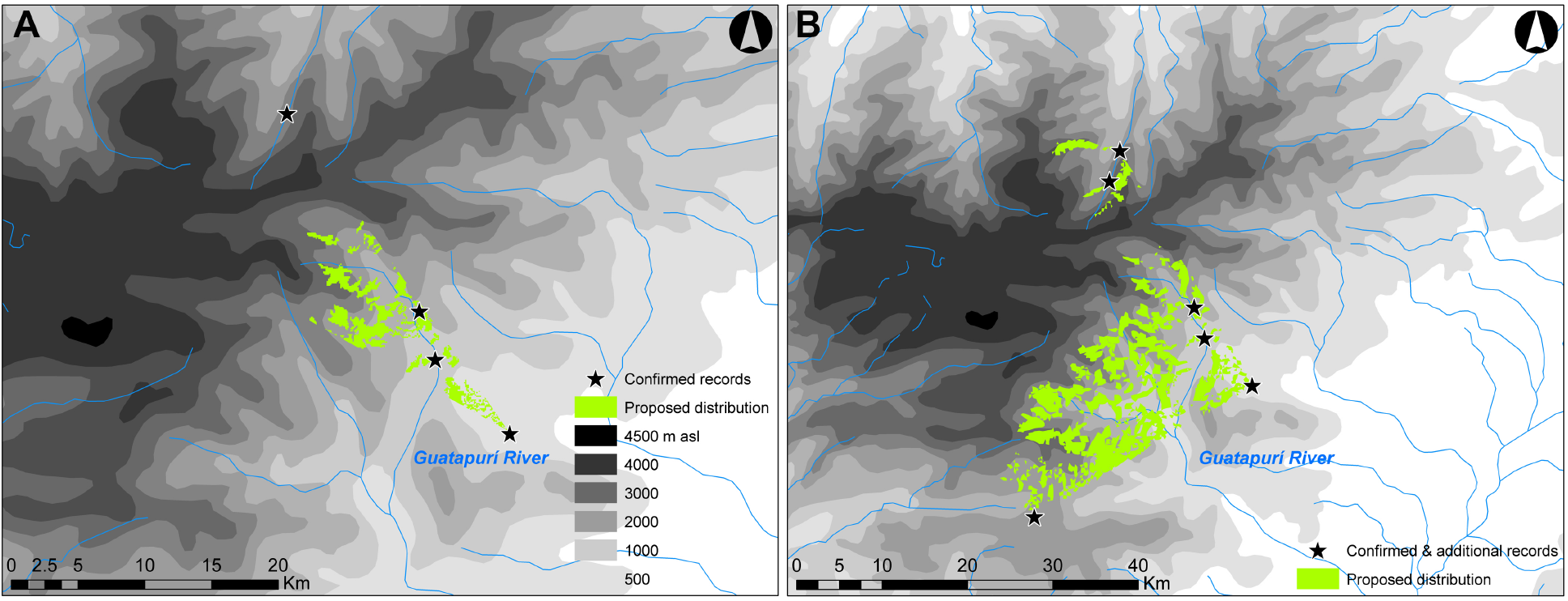
Proposed area of occupancy (AOO) of Santa Marta Sabrewing delimited by the extension of suitable habitats (excluding the 2010 San Lorenzo record, see text). Distribution hypotheses consider (A) only confirmed records of the species, and (B) confirmed records and all observations on the southern and north-eastern slopes of the SNSM described by F. A. A. Simons during his expedition (Salvin and Godman 1879).

### Local occupancy rate and seasonal movements

We detected Santa Marta Sabrewing in nine out of 20 point-count stations surveyed, resulting in a naive occupancy estimate (*sD/s*) of 0.45. The intercept-only model indicated that the species’ occupancy rate and detection probability were 0.46 (standard error SE = 0.11) and 0.63 (0.09), respectively (see Supplementary Material Table S2). The “saturated” occupancy model was not conclusive regarding a potential effect of elevation on Santa Marta Sabrewing occupancy (Supplementary Material Table S2). The “elevation-only” model fitted the data better and suggested a very small increase in occupancy with elevation (Supplementary Material Figure S1), although this effect had weak statistical support (Supplementary Material Table S2).

The raw abundance of Santa Marta Sabrewing was similar between the lower (1,150–1,550 m asl) and higher elevational ranges (1,550–1,900 m) of the study area, with an average of 0.39 and 0.44 birds/count. While only 0.2 birds/count were recorded in stations that were mainly surrounded by transformed areas, 0.6 birds/count were detected in stations where land cover was dominated by native vegetation (i.e., secondary vegetation and early growth, as defined previously). The highest raw abundance corresponded to point count stations surrounded mostly by native vegetation below 1,550 m, with an average of 1.4 birds/count.

We confirmed the presence of Santa Marta Sabrewing at the rediscovery site (∼1,150– 1,850 m) for 16 consecutive months between July 2022 and October 2023. Males and a few females were recorded during every visit, with males always detected in the same spots. We found no evidence of birds migrating.

### Habitat associations, foraging, and behaviour

Of the 23 sabrewing point-count records, 78% were in secondary vegetation and 22% in mixed vegetation. No records were in point-counts in early growth areas or open pastures. Neu’s habitat selection index suggested that secondary vegetation was preferred (1.2) to the remaining habitats (early growth: 0; mixed: 0.9; open: 0). However, neither habitat was used more frequently than expected given its areal extent (Fisher’s exact test *P* = 0.74). We obtained 25 behavioural records from *ad libitum* observations of the sabrewing. Away from the point counts, sabrewings were mainly observed in remnant riparian forest strips, all of which had been heavily disturbed, on steep slopes along La Macana stream and directly at the Guatapurí River above Guatapurí village. Individual hummingbirds were observed visiting agroforestry systems from nearby patches of secondary vegetation and early growth. We grouped all sabrewing observations into three basic behavioural categories: foraging (12% of records); flying (36%); and vocalizing in perches (52%). Most foraging observations were obtained in mixed habitat, particularly in agroforestry systems with shade-grown coffee, banana, and sugarcane. Females and males took nectar from flowering banana trees (*Musa* sp.) and from 7–14 m tall *Calliandra* and *Erythrina* trees. One foraging event involved a male taking nectar from the flowers of *Malvaviscus concinnus* 1.7 m above the ground in a riparian forest. Most observations of “flying” and “vocalizing in perches” involved males, which vocalized non-stop during the morning (∼07:30–12:00) while defending their territories.

Males would defend territories alone or aggregated in loose leks composed of up to 4 individuals located ∼9–15 m apart (*n* = 2 leks). Two leks were found inside 30–60 m wide and 12– 18 m tall patches of riparian forest at elevations of 1,551 and 1,746 m, respectively. Territorial behaviour was observed year-round. Males used perches 3–7 m above the ground to vocalize throughout the morning and defended their territory from conspecifics and other hummingbird species by aggressively chasing intruders in flight (Supplementary Material Appendix 2). We also observed displays that consisted of males slowly approaching an intruder with a threatening posture, hovering forward with tail feathers and wings extended to appear larger (Supplementary Material Appendix 2). In November 2022, a different display-like manoeuvre was observed: Two males engaged in a vertical ascent display (15–20 m), after which they turned around and performed a display dive with their tail feathers open.

We obtained only one observation that could indirectly suggest breeding activity in the population of sabrewings at La Macana stream. This was a female collecting spider web with her bill in May 2023 that could be an individual gathering material for nest building.

### Vocalizations

The Santa Marta Sabrewing male territorial vocalizations we recorded are composed of single (one-type) notes of short duration. The number of notes produced per minute (i.e., calling rate) was variable and may be associated with conspecific local density (e.g., isolated vs. lekking males). Male A, which was observed alone, had the lowest calling rate (42 notes/min), whereas Male B that was observed at a “crowded” lek site had the highest calling rate (93 notes/min), and Male C had an intermediate calling rate (78 notes/min; see Figure 3). The mean note duration was relatively similar among males (Male A: 0.024 s; B: 0.026 s; C: 0.036), but the duration of silent intervals showed more variation (Male A: 1.177 s; B: 0.608 s; C: 0.696). The mean spectral frequencies were 4.257 kHz (SD = 0.459 kHz) for Male A, 3.570 kHz (0.385) for Male B, and 3.577 kHz (0.437) for Male C. Peak of frequency spectrum were 4.50, 3.78, and 3.87 kHz for males A, B, and C, respectively. Male A, which had the lowest calling rate, had the highest mean and peak frequencies (Figure 4).

**Figure 4.**
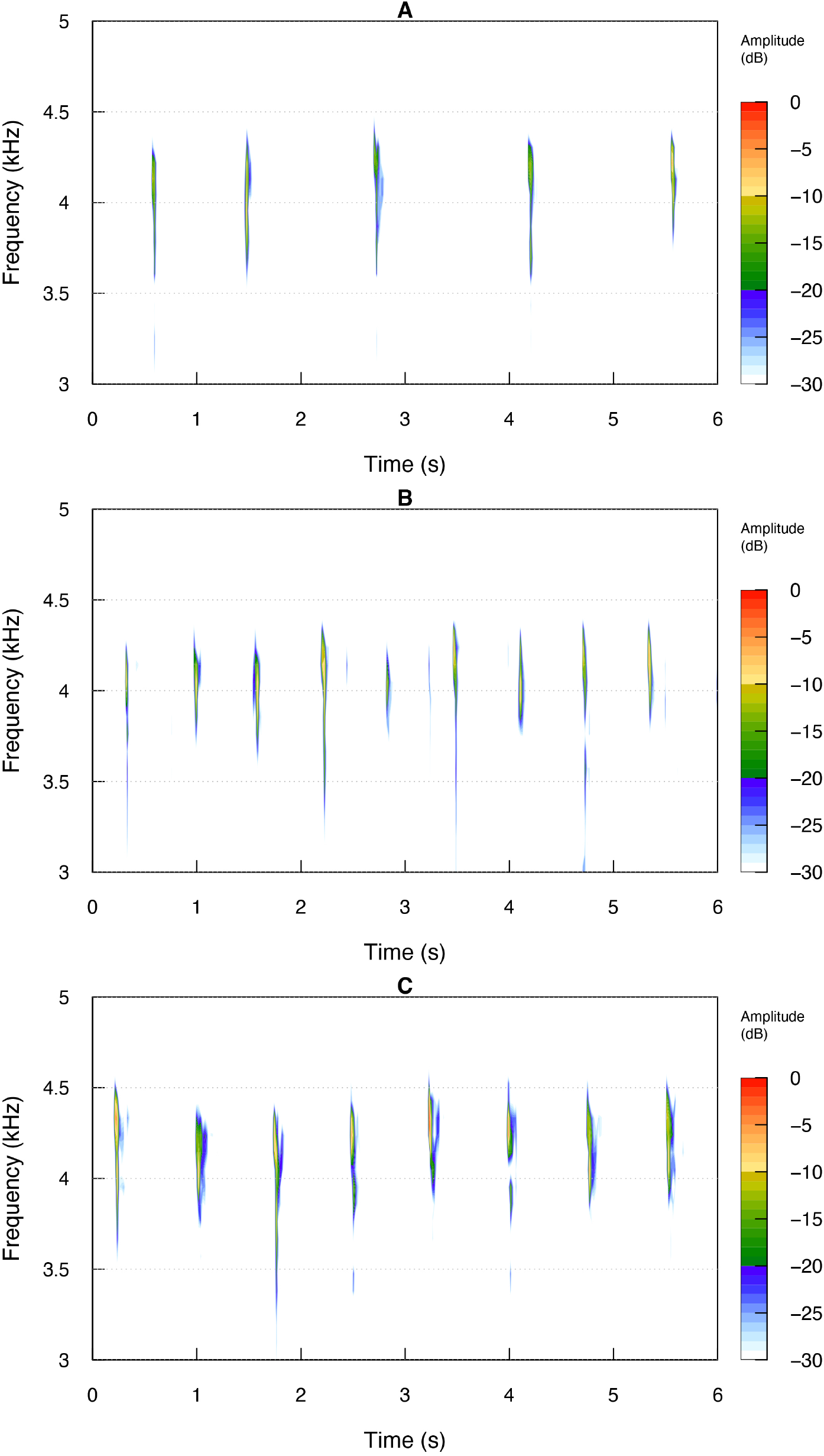
Sonograms of Santa Marta Sabrewing’s calls for three different individuals (males A, B, and C; see main text). The horizontal axis indicates time (in seconds), and the vertical axis indicates frequency (in kHz). Audio files for the three males are deposited at the Macaulay Library scientific archive (https://macaulaylibrary.org) with catalog numbers ML615927194, ML615917198, and ML615927203.

## Discussion

This is the first study to combine historical and reliable recent information on Santa Marta Sabrewing to describe aspects of its distribution and natural history, while contributing to an increased understanding of its conservation status. From a conservation perspecitve, our results support the assesment of Santa Marta Sabrewing as CR and highlight the need for more information to understand its distribution, habitat preferences, and population size. Given the high number of erroneous and unsupported sabrewing records that we uncovered during our study, we argue that it is essential for new records of Santa Marta Sabrewing to be rigorously documented and verified to avoid false positives resulting from misidentifications. This particularly applies to records for previously unreported localities. Our study shows that a local population of Santa Marta Sabrewing can be found throughout the year in an area that coincides with the type locality and nearby areas where the species was first observed by Simons almost 150 years ago.

### Historical distribution and seasonal movements

The assumption that Santa Marta Sabrewing had a wide distributional range, with individuals commonly observed up to the snowline, seems to originate from Salvin and Godman’s (1879) transcriptions of Simons’ comments during his eight-month journey across the SNSM in 1878 (Simons 1879, 1881). In his notes, Simons mentions that Santa Marta Sabrewing was found as high as ∼4,500 m, and stated that sabrewings were commonly observed in June and July at ∼3,300 m, in addition to being found at lower elevations in San Miguel (La Guajira Department) at ∼1,800 m and in San Sebastián de Rábago (Cesar Department) at ∼2,050 m (Salvin and Godman 1879). In contrast to these comments, however, most of Simons’ specimes were collected at elevations below 2,500 m (Salvin and Godman 1879), and neither the sabrewing specimens he collected nor the ones collected by Carriker in 1945 and 1946 came from localities above 2,800–3,000 m. Apart from these notes by Simons, no one else has observed the sabrewing at higher elevations. Thus, it is difficult to establish with certainty whether the species was indeed present at higher elevations, as has been suggested for decades. Simons was helped by indigenous people while crossing the snowline via the headwaters of the Guatapurí River (> 4,000 m), and there is a chance that some of his observations of Santa Marta Sabrewing occurred during that journey. Revisiting this area would help resolve this uncertainty.

Similar to the assumptions about distribution, the supposition that Santa Marta Sabrewing is an elevational migrant is based solely on Simons’ notes (see Salvin and Goodman 1879). Our review found no empirical data that corroborate Simons’ idea and in our fieldwork we observed Santa Marta Sabrewings, likely the same individuals, over 16 consecutive months at La Macana stream, including two wet seasons. Considering the current knowledge on the ecology of *Campylopterus* sabrewings (McMullan 2016, Ayerbe Quiñones 2018, Winkler et al. 2020) and our data, we propose three hypotheses regarding the distribution and possible migratory movements of the species. First, the sabrewing is a partial migrant with a portion of the population seasonally migrating to the paramo, while the remaining individuals occupy their territories at mid-elevation habitats year-round. Second, the species is a facultative migrant whose migrations depend on inter- and intra-annual fluctuations in precipitation and resource availability. Third, the species is a resident that does not migrate, and records outside the core area of distribution in the Guatapurí basin are vagrants.

While the data remain sparse, our results favour the third hypothesis. In addition to the continuous presence of the species at La Macana stream, the photograph from San Lorenzo IBA is the only documented record of Santa Marta Sabrewing outside of the Guatapurí basin. The San Lorenzo record has been widely accepted by the ornithological community for more than a decade, but in our review it emerged as a clear outlier relative to the other confirmed records both in its geographic location and in its pattern of detection. This, combined with the fact that the best-known fieldmark for distinguishing between Lazuline and Santa Marta Sabrewing (the bird’s tail) is not fully visible in the photograph, suggest that it could be worth reexamining this photograph as our knowledge of the identification of Santa Marta Sabrewing continues to improve. Regardless of how the San Lorenzo record is evaluated, however, it seems very unlikely that it represents a resident population or a consistent pattern of migration. There have been extensive ornithological surveys and birding effort in San Lorenzo before and since the observation and our data, like Simons’ and Carriker’s, suggest that Santa Marta Sabrewings have a relatively high detectability when present (see below). Instead, Santa Marta Sabrewing appears to be a case of microendemism, probably restricted to the south-eastern slope of the SNSM, and specifically the Guatapurí basin (Figure 3). Futher research may reveal the species to occur in other parts of the SNSM, such as remnant forests of the northern flank (e.g., San Miguel and Rioancho, La Guajira Department), or west of the Guatapurí River (e.g., San Sebastian de Rábago; Figure 3B), but our results indicate that the burden of proof should be on confirming the species’ speculated migratory behavior and presence outside of the Guatapurí basin rather than vice versa.

### Natural history and habitat associations

We found sabrewings to be highly localized in the rediscovery site, but conspicuous within this area. The species was detected with a probability of 0.6 in our surveys at the site and the birds vocalized repetitively during the morning, year-round. Simons also commented on sabrewings being common around Atánquez, and that their presence could be noticed with ease when vocalizing (Salvin and Godman 1879). The fact that the species can be reliably detected when present points to very low numbers and a limited distribution, rather than difficulty of detection, being the reason that it has been reported so rarely.

Our observations and those of Simons suggest that this species is more abundant in mid-elevation forests, possibly in the vicinity of watercourses. We found no quantitative evidence of Santa Marta Sabrewing selecting forests over disturbed habitats, but the presence of territories and leks in riparian strips point to forests as an important habitat. Although the species uses cultivated areas, we hypothesize that its presence in transformed landscapes might be conditioned by the presence of native vegetation in association with watercourses. Thus far, we have found leks in dense understorey in riverine forest strips, and the presence of riverbanks near leks may indicate that female nesting sites could be nearby (see Esteves Lopes et al. 2020). Nest sites in the vicinity of streams has been reported in other *Campylopterus* species (Hayes et al. 2000, Marín 2001, Hayes 2002).

Information on Santa Marta Sabrewing diet and breeding phenology remains extremely limited. From our review and field observations, the species has been seen consuming nectar from plants of five families, including Ericaceae (*Befaria*), Fabaceae (*Calliandra, Erythrina*, and *Inga*), Malvaceae (*Malvaviscus*), Musaceae (*Musa*), and Rubiaceae (*Palicourea*). This list can help inform conservation actions aimed at increasing suitable habitats in productive landscapes. Regarding the species’ breeding phenology, our only observation of what could be breeding behavior occurred in May, coinciding with reports of courtship activity during April–June (reviewed in Cárdenas-Ortiz and Cortés-Herrera 2016).

### Conservation, threats, and future work

The EOO projected in this study was much lower than the values reported in previous extinction risk assessments (2,900 km^2^ in BirdLife International 2023b; 7,702 km^2^ in Renjifo et al. 2016). Similarly, our AOO projection was lower than the inferred remnant habitat (4,004 km2; Renjifo et al. 2016). If a RedList category were to be applied based on criteria B1 and B2 and the estimates provided here, the species would be classified as endangered (EN). This is in line with the latest extinction risk assessment carried out by BirdLife International, which lists Santa Marta Sabrewing as endangered (EN) based on criterion B1 and CR based on criterion C2 (inferred small and declining population size). If the San Lorenzo record is not included in the model, the species’ AOO might only cover an area of ∼23 km^2^, if restricted to the Guatapurí basin (Figure 3A), or at most ∼164 km^2^ if the other localities mentioned by Simons are included (Figure 3B). Furthermore, if more conservative EOO and AOO projections are made by considering all documented historical presences, the species should remain listed as EN based on criteria B1 and B2 (∼1,571 and ∼563 km^2^, respectively; Supplementary Material Figure S2). We thus recommend that the species remain listed as CR until conclusive evidence suggests otherwise.

Just a few years before its rediscovery in 2022, Santa Marta Sabrewing was feared to have become extinct in the wild (Salaman et al. 2022). We have found what seems to be a stable but likely small population along La Macana stream in Chemesquemena, and this raises hope for the survival of the species. Most of the species’ hypothesized distribution lies within the Sierra Nevada National Park (Supplementary Material Figure S3), and the national government recently agreed with four indigenous groups on a long-term management plan and the expansion of the park. A strategy to strengthen governance is required to accompany this new arrangement given that the protection of natural ecosystems within the park has not been effective (Duran-Izquierdo and Olivero-Verbel 2021). The focal population near Chemesquemena and at San José is currently unprotected, and despite being located within Kankuamo and Kogi indigenous reserves, land-use practices (e.g., fires and forest clearing) in the Guatapurí basin may pose a threat. Sugarcane cultivation by indigenous peoples has been documented since the 19th century (Salvin and Godman 1879), and is still widely practiced across the region. This practice usually leads to deforestation of strips of native riparian vegetation, which buffer river floods and protect local family lands and other crops. Using agroforestry systems instead of sugarcane can help conserve the population of Santa Marta Sabrewing along La Macana stream and ensure environmental sustainability and resilience of local peoples’ agricultural productivity.

Santa Marta Sabrewing has been labeled as perhaps the rarest and most threatened bird species in Colombia (Salaman et al. 2022), and continued studies are urgently needed to design effective conservation action for the species. First, further field surveys are necessary to clarify the distribution and produce records so that distribution models can be used to form a realistic distribution hypothesis for Santa Marta Sabrewing. Once it is feasible to use distribution models, other relevant measurements (e.g., EOO and AOO) will be available to better assess the species’ risk of extinction. Second, conducting a longer and more widespread study of the population at La Macana stream is necessary to determine whether Santa Marta Sabrewing occupancy and abundance vary with elevation and other relevant environmental factors. This will also make it possible to assess whether individuals we monitored remained at mid-elevations during our study given the constantly humid conditions due to the past La Niña event, which spanned approximately three years (2020–2023). Finally, further efforts are needed to conduct nest searches during April– June along La Macana stream and its tributaries to characterize the species’ habitat requirements during this sensitive period of the annual cycle.We hope that the characterization of the Santa Marta Sabrewing vocal repertoire and behavioral displays will facilitate the correct identification of this species, and in doing so help to increase information on its biology, ecology, and distribution.

The sabrewing populations we studied along La Macana stream occur inside a private reserve of the Kankuamo indigenous community. Any further study and observation requires the active participation of the indigenous landowners and collaboration in developing conservation measures. We extend this call to the general public and the actors involved in the birdwatching tourism industry, who can help the local community develop a sustainable ecotourism strategy that benefits the Kankuamo people and the region’s endemic and threatened avifauna.

## Acknowledgements

We are thankful to all those who allowed us to visit the study area, particularly the Cabildo Indígena del Resguardo Kankuamo and the Chemesquemena community, as well as the Kogi people of San José. We also thank Luz Eneida Ochoa and Jaider Carrillo for their support and Elquin Toro for his valuable help in the field and for facilitating photos used in this manuscript. Special thanks to Cristobal Navarro for his advice and Gustavo Bravo for checking specimens at the IAvH Ornithological Collection. We also thank Andrés Gómez, Leonardo Lemus, María Camila Machado, Gina Gómez Junco, and Daniel J. Lebbin. This work was funded by American Bird Conservancy and the Search for Lost Birds (grant No. 20154F), World Parrot Trust, SELVA, ProCAT Colombia, and Universidad Nacional de Colombia.

No birds were captured or manipulated during this study. Fieldwork protocols were approved under Resolution No. 179-2015 from Parques Nacionales Naturales de Colombia and Resolution 0597-2014 and 0189-2016 from Autoridad Nacional de Licencias Ambientales issued to SELVA.

Data analyzed in this study and supplementary media files are available from the Zenodo open data repository at https://doi.org/10.5281/zenodo.10651684.

