## Supplementary Material for "Distribution, ecology, and natural history of the recently rediscovered and critically endangered Santa Marta Sabrewing"

[1]SELVA: Research for Conservation in the Neotropics, Bogotá, Colombia.
[2]Universidad Nacional, sede de La Paz, Cesar, Colombia.
[3]World Parrot Trust
[4]Proyecto de Conservación de Aguas y Tierras – ProCAT Colombia
[5]Instituto de Ciencias Naturales, Universidad Nacional de Colombia, Bogotá, Colombia
[6]American Bird Conservancy, P.O. Box 249, The Plains, VA 20198, USA

### Appendix 1: supplementary tables and figures

**Supplementary Table S1**
Historical records of Santa Marta Sabrewing in the Sierra Nevada de Santa Marta, northern Colombia, with documented evidence in the form of specimens or photographs. Validity of the San Lorenzo record needs further examination.

| Locality | Department | Latitude | Longitude | Type | Source |
|---|---|---|---|---|---|
| San José | Cesar | 10.75 | -73.4 | Preserved specimen | GBIF |
| Atánquez | Cesar | 10.7 | -73.35 | Preserved specimen | GBIF |
| Chenducua | Cesar | 10.78 | -73.41 | Preserved specimen | GBIF |
| Alto de Macotama | Guajira | 10.92 | -73.50 | Preserved specimen | GBIF |
| San Lorenzo** | Magdalena | 11.10 | -74.07 | Photograph | eBird |

**Supplementary Table S2**
Candidate model set describing variation in the local occupancy rate of Santa Marta Sabrewing in a recently rediscovered population along La Macana stream, Sierra Nevada de Santa Marta, northern Colombia. Models estimated occupancy either as a constant or as a function of elevation and the proportion of native vegetation within point-count stations. Detection probability was held constant for all models. Logistic-linear parameter estimates (±SE) and Akaike information criterion (AIC) values are shown for each model. All predictors were standardized prior to model fitting.

| **Model parameter** | **Estimate** | **SE** | ***z*** | ***P*** |
|---|---|---|---|---|
| *Intercept-only model (AIC = 78.29)* | | | | |
| Intercept | -0.17 | 0.46 | -0.36 | 0.72 |
| Detection probability | 0.52 | 0.37 | 1.41 | 0.16 |
| *Elevation-only model (AIC = 72.38)* | | | | |
| Intercept | 1.56 | 1.35 | 1.15 | 0.25 |
| Elevation | 0.98 | 1.66 | 0.59 | 0.55 |
| Elevation$^2$ | -3.16 | 2.12 | -1.48 | 0.14 |
| Detection probability | 0.51 | 0.37 | 1.40 | 0.16 |
| *Saturated model (AIC = 74.11; ĉ = 0.81)** | | | | |
| Intercept | 2.17 | 2.06 | 1.05 | 0.29 |
| Elevation | 2.05 | 2.89 | 0.71 | 0.48 |
| Elevation$^2$ | -3.65 | 2.59 | -1.41 | 0.16 |
| Proportion of native vegetation | -1.73 | 3.67 | -0.47 | 0.64 |
| Detection probability | 0.49 | 0.38 | 1.31 | 0.19 |

* ĉ = 0.81; chi-square goodness-of-fit test $\chi^2$ = 7.49, *P* = 0.8.

**Supplementary Figure S1**

Variation in the local occupancy rate of Santa Marta Sabrewing as a function of elevation in a recently rediscovered population along La Macana stream, Sierra Nevada de Santa Marta, northern Colombia. Predictions were generated from a single-season occupancy model that included the linear and quadratic effects of elevation as predictors, and held detection probability constant.

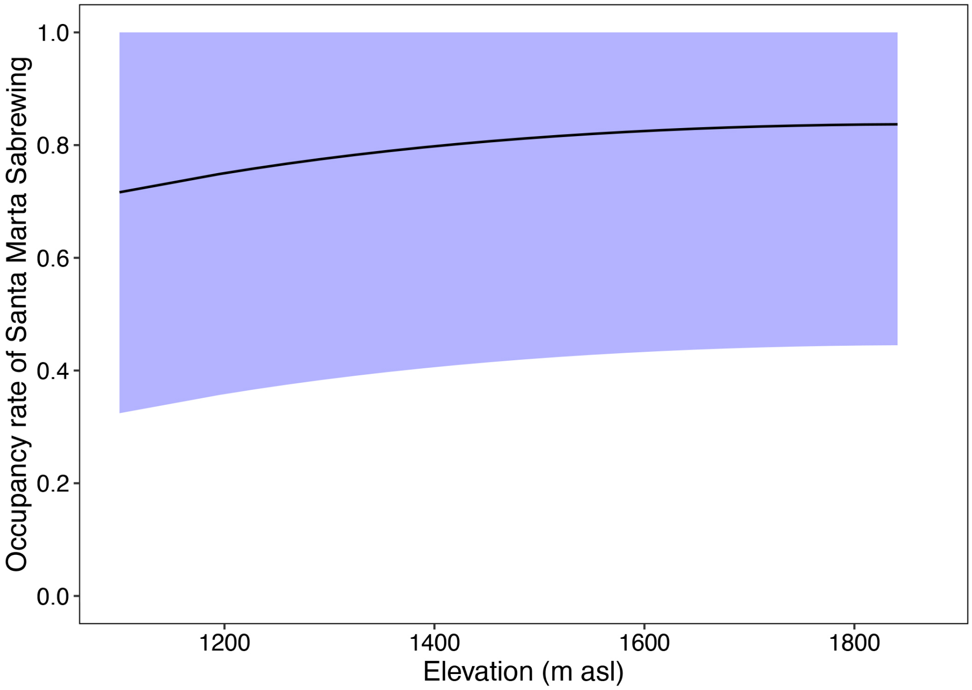


**Supplementary Figure S2**

Spatial representation of Santa Marta Sabrewing's (A) extent of occurrence (EOO), and (B) area of occupancy (AOO) in the Sierra Nevada de Santa Marta, northern Colombia, considering all documented historical records. This includes observations on the southern and north-eastern slopes of the SNSM described by F. A. A. Simons during his expedition (Simons 1879, 1881; see also Salvin and Godman 1879).

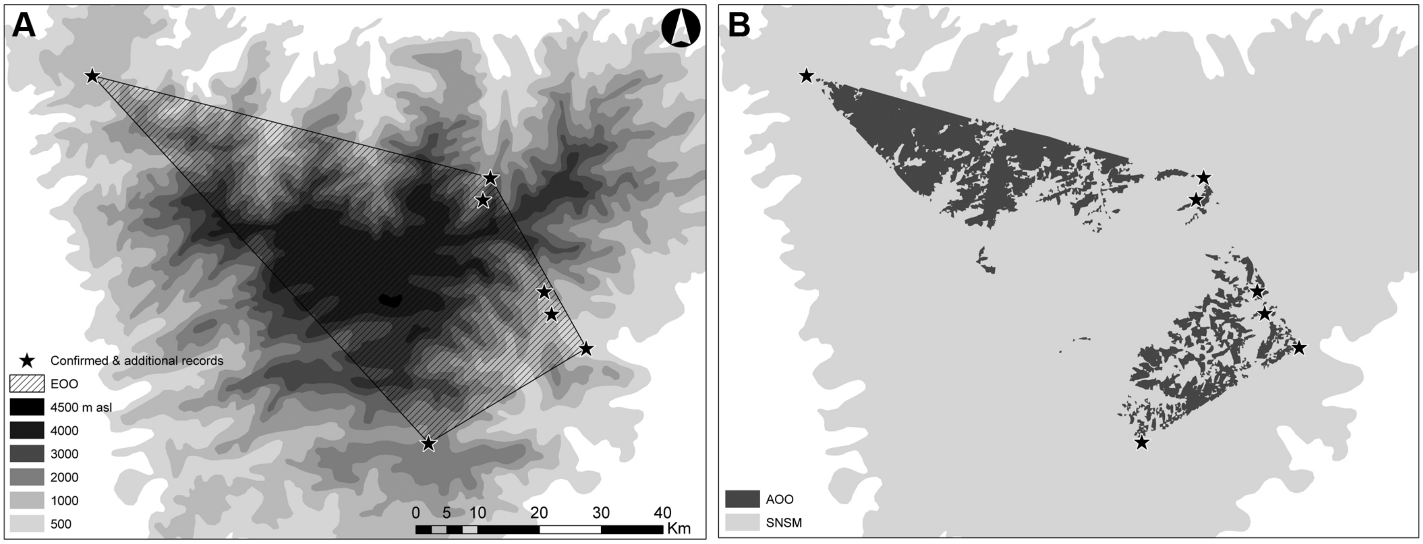

**Supplementary Figure S3**
Overlapping of Santa Marta Sabrewing area of occupancy (AOO) with the Sierra Nevada de Santa Marta National Park. The species' AOO is the same shown in Supplementary Figure S2.

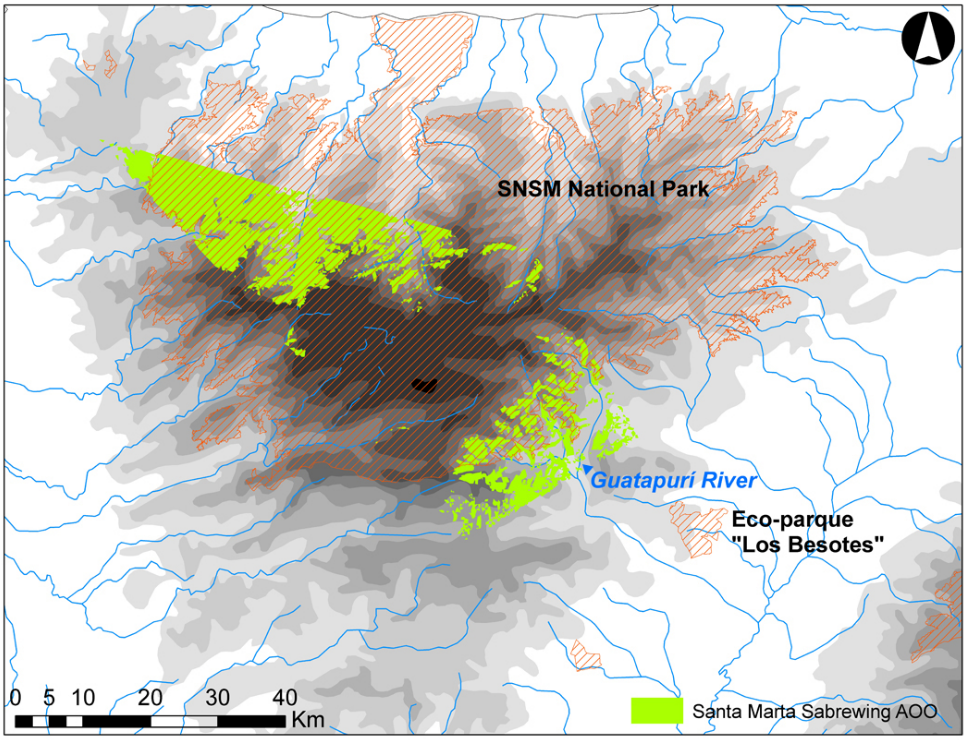

### Appendix 2: supplementary files

This supplement is accompanied by media files that support behavioural descriptions in the main text (see Botero-Delgadillo et al. 2024). No exact coordinates from the focal population were included in this supplement or related documents/materials to protect these locations.

**Supplementary Video S1**
A lek-attending male Santa Marta Sabrewing vocalizing in a perch and subsequently adopting a threatening body posture with wing and tail feathers extended towards a conspecific male intruder. Footage was taken along the course of La Macana stream, near Chemesquemena village (Cesar Department, northern Colombia). The "Campylopterus_threat.mp4" video accompanies this supplement. Video: Elquin Toro © (reproduced with permission).

**Supplementary Video S1**
A slow-motion (~0.1 x) video showing a perching male Santa Marta Sabrewing involved in a persecution flight with a conspecific male aggressor. Footage was taken along the course of La Macana stream, near Chemesquemena village (Cesar Department, northern Colombia). The "Campylopterus_fight.mp4" video accompanies this supplement. Video: Elquin Toro © (reproduced with permission).

Salvin, O., and Godman, F. D. C. (1879). On a Collection of Birds from the Sierra Nevada of Santa Marta, Columbia. *Ibis* **21:** 196–206.

Simons, F. A. A. (1879). Notes on the Topography of the Sierra Nevada of Santa Marta, U. S. of Colombia. *Proc. Roy. Geogr. Soc. & Monthly Rec. Geogr.* **1:** 689–694.

Simons, F. A. A. (1881). On the Sierra Nevada of Santa Marta and its Watershed (State of Magdalena, U. S. of Colombia). *Proc. Roy. Geogr. Soc. & Monthly Rec. Geogr.* **1:** 705–723.
